# Neurophysiological Correlates of the Rubber Hand Illusion in Evoked and Oscillatory Brain Activity

**DOI:** 10.1101/136846

**Authors:** Isa S. Rao, Christoph Kayser

## Abstract

The rubber hand illusion (RHI) allows insights into how the brain resolves conflicting multisensory information regarding body position and ownership. Previous neuroimaging studies have reported a variety of neurophysiological correlates of illusory hand ownership, with conflicting results likely originating from differences in experimental parameters and control conditions. Here, we overcome these limitations by using a fully automated and precisely-timed visuo-tactile stimulation setup to record evoked responses and oscillatory responses in the human EEG. Importantly, we relied on a combination of experimental conditions to rule out confounds of attention, body-stimulus position and stimulus duration, and on the combination of two control conditions to identify neurophysiological correlates of illusory hand ownership. In two separate experiments we observed a consistent illusion-related attenuation of ERPs around 330 ms over frontocentral electrodes, as well as decreases of frontal alpha and beta power during the illusion that could not be attributed to changes in attention, body-stimulus position or stimulus duration. Our results reveal neural correlates of illusory hand ownership in late and likely higher-order rather than early sensory processes, and support a role of premotor and possibly intraparietal areas in mediating illusory body ownership.

## 1 Introduction

Philosophy, psychology, and neuroscience continue to debate the sources and modulators of conscious experience. The scientific study of consciousness has long been focussed on the visual domain, but recent decades have seen a rise of interest in bodily self-consciousness and the integration of bodily signals with other multisensory information (Jeannerod, 2007). Bodily self-consciousness refers to the integrated, pre-reflexive experience of being a self in a body and has been related to tactile, vestibular, proprioceptive, as well as visual and motor information (Blanke, 2012; Tsakiris and Haggard, 2005). One extensively investigated aspect of bodily self-consciousness is the experience that our body and its parts belong to us and are distinguished from non-body objects and other people’s bodies, so-called body ownership. A widely used paradigm to study body ownership is the rubber hand illusion (RHI; Botvinick, 2004) during which participants watch an artificial rubber hand being stroked in synchrony with strokes on their own occluded hand. This synchronous visuo-tactile stimulation alters bodily experience as it induces the illusion that the rubber hand is one's own hand.

Several functional magnetic resonance imaging (*fMRI*) studies have aimed to identify the neural correlates of illusory hand ownership. The experience of illusory hand ownership has been linked to activity in frontal brain regions, such as the premotor cortex (Bekrater-Bodmann et al., 2014; Ehrsson, 2004; Petkova et al., 2011), occipito-temporal regions such as the extrastriate body area (Limanowski et al., 2014), intraparietal areas (Petkova et al., 2011), the anterior insula (Limanowski et al., 2014), and the temporoparietal junction (Guterstam et al., 2013). However, given the nature of the fMRI signal, these studies have not been able to provide a functionally specific picture that assigns these neural correlates to a specific part of the sensory-perceptual cascade, for example by assigning the relevant neural activations to a specific latency following each repeat of the visuo-tactile stimulation.

Overcoming these limitations, several electroencephalographic(EEG) studies have aimed to reveal the physiological correlates of illusory hand ownership at higher temporal precision. One such study has described the relative attenuation of somatosensory-evoked responses during the Illusion about 55 ms after stimulus onset (Zeller et al., 2015). This attenuation was localized to the primary somatosensory cortex and the anterior intraparietal sulcus, and was interpreted in the context of predictive coding models as an attenuated precision of the relevant proprioceptive representations that are required to solve the multisensory conflict induced by the RHI. However, another EEG study using a similar experimental paradigm reported illusion-related changes in ERPs only at much longer latencies of around 460 ms over frontal electrodes (Peled et al., 2003). As a result it remains unclear whether neural correlates of the RHI include aspects of early sensory encoding, hence at shorter latencies relative to stimulus onset, or mostly involve higher cognitive processes emerging at longer latencies relative to the touch stimulus.

The lack of clear insights from the existing EEG studies on the RHI may in part result from the use of different stimulation parameters and the use of distinct control conditions. Two widely used control conditions for the rubber hand illusion are the Incongruent condition, in which the rubber hand is placed as an anatomically incongruent angle, and the Real condition, in which the rubber hand is absent and stimulation occurs on the real hand in view (Ehrsson, 2004; Olivé et al., 2015; Schmalzl et al., 2014; Tsakiris et al., 2007; Zeller et al., 2015, 2016). Unfortunately, these control conditions carry inherent confounds by differing from the illusion condition by more than just the absence of the illusion. In the Real condition, the hand position is changed and the rubber hand is completely absent from the setup, hence all seen potential body parts are indeed a natural part of the participant’s body. In the Incongruent condition the visual stimulation of the Rubber Hand and the somatosensory stimulation on the real hand occur in two different locations, while in the Illusion condition these stimulations are perceived to occur on one location, i.e. on the rubber hand. It is hence possible that spatial attention in the Illusion condition is focused on one location, while in the Incongruent condition attention is divided across two locations. In addition, in the Illusion condition, the visual stimulus is perceived to occur on the participant’s body, i.e. the embodied rubber hand. The visual stimulus in the Incongruent condition however is perceived to occur not on the body, but on the non-embodied rubber hand. Previous studies have suggested that visual stimuli are processed differently when the stimuli is placed near the hand, rather than when it is not (Langerak et al., 2013). Thus, visual processing due to body-stimulus position between Illusion and Incongruent condition may differ substantially. As a result it remains unclear whether illusion-related effects reported in previous studies are indeed only related to the illusory body experience, or rather originate from other confounding factors in the experimental paradigm. Hence, it remains very difficult to collate findings across studies and to reliably identify the electrophysiological correlates of illusionary hand ownership.

Our goal was to study the neural correlates of the RHI in electroencephalographic brain activity by refining the typical protocol used to induce the RHI in two ways: First, by introducing a temporally precise stimulation apparatus that allows the recording of evoked activity that is precisely-time locked to the somatosensory stimulus; and second, by comparing neural correlates of the RHI across different control conditions to rule out confounds of attention and body-stimulus position. Given that previous studies have reported illusion-related effects both in evoked responses (Peled et al., 2003; Zeller et al., 2015) and in induced oscillatory activity (Faivre et al., 2017), we here focused on both these markers of neural processing. In the first experiment we recorded EEG activity during the Illusion, the Real and Incongruent control conditions and two further conditions which varied in attention focus and body-stimulus position. We identified neurophysiological correlates of illusionary hand ownership that were consistent across both control conditions and then differentiated these from the two confounds. In a second experiment, we replicated these neurophysiological correlates of illusionary hand ownership, and furthermore established that these are independent of stimulus duration.

## 2 Materials and Methods

### 2.1 Participants

Forty right-handed volunteers (Experiment 1: n=20 participants including 13 female, mean age = 23.1 years, SD = 3.1; Experiment 2: n=20 participants including 13 female, mean age = 22.1 years, SD = 2.9 years) participated in the study. All participants had taken part in a previous experiment involving the rubber hand illusion. Only participants who agreed or strongly agreed to the statement ‘During the last trial I felt as if the rubber hand were my hand’ (Botvinick and Cohen, 1998) during this previous experiment were invited to take part in the current study. All participants gave written informed consent before participation in this study. All protocols conducted in this study were approved by the Ethics Committee of the College of Science and Engineering of the University of Glasgow.

### 2.2 Experimental conditions

Participants sat on a comfortable chair in front of a one-compartment, open-ended box placed on a two-storey wooden platform. Their left arm was placed on an arm rest. Visual stimulation was delivered by a red light-emitting diode (LED; Seeedstudio, 10mm diameter) positioned 5 cm to the right of the box on the top storey. Tactile stimulation was delivered by a vibration motor placed close to the subject’s skin (Permanent magnet coreless DC motor, Seeedstudio, 10mm diameter). Visual and tactile stimulation were controlled via Matlab and an Arduino prototyping platform.

In experiment 1 five conditions were administered in a randomised order (Fig. 1). Illusion: The participant’s left hand was placed in the box with the tip of the middle finger positioned on a vibration motor. The right hand was placed at the other end of the platform in reaching distance of the keyboard. A lifelike rubber hand was positioned in an anatomically congruent orientation next to the box in a distance of 15cm to the participant’s hidden left hand. The middle finger of the rubber hand was placed on a dummy vibration motor. The LED was positioned five millimetres above the dummy motor. This condition is typically used to induce the RHI. Incongruent: The rubber hand was placed at an angle of 45°. Besides this the setup was similar to the setup described in Illusion (Ehrsson, 2004; Olivé et al., 2015; Press et al., 2008; Zeller et al., 2015, 2016). Real: No rubber hand was present. The middle finger of the left hand was placed in view on a vibration motor positioned 5 millimeters below the LED. The right hand was in the same position as in the Illusion and Incongruent conditions (Zeller et al., 2015, 2016). Hand under: The participant’s left hand was placed on the lower storey of the platform with the middle finger placed on a vibration motor. The vibration motor was positioned right below the dummy vibration motor on the top storey. The vertical distance between the two motors was 10 cm. Besides this, the setting was identical to the Incongruent condition. Two hands: No rubber hand was present. The middle finger of the participant’s right hand was placed on a dummy vibration motor below the LED. Besides this, the setting was identical to the Incongruent condition. Throughout all conditions view of the left arm, and the trunk of the rubber hand where applicable, was obstructed by an opaque piece of fabric.

**Figure 1.**
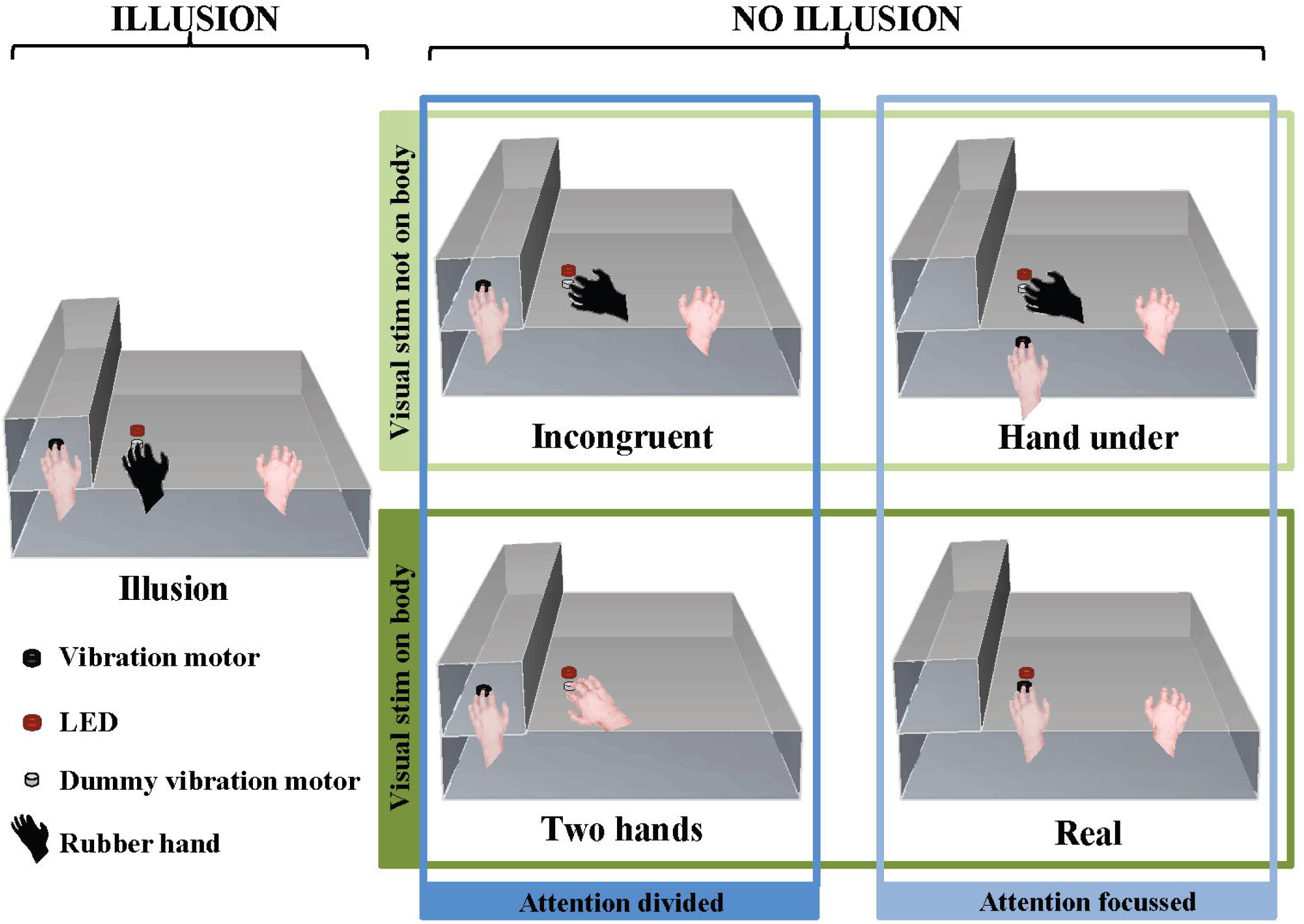
Experimental setup during the five conditions. Illusion: Congruently placed rubber hand on dummy vibration motor below LED, left hand on vibration motor. Incongruent: Incongruently placed rubber hand on dummy vibration motor below LED, left hand on vibration motor. Hand under: Incongruently placed rubber hand on dummy vibration motor below LED, left hand on vibration motor below dummy vibration motor and LED. Two hands: No rubber hand, left hand on vibration motor, right hand on dummy vibration below LED. Real: No rubber hand, left hand on vibration motor under LED. The four non-illusion conditions were additionally grouped in a 2x2 design according to the factors attention and body-stimulus position.

The differences in hand and stimuli location across conditions allow for a grouping of Incongruent, Real, Hand under and Two hands in regards to attentional and body-stimulus related processing (Fig. 1). In the Incongruent and Two hands conditions attention is divided, since in both conditions visual and somatosensory stimuli occur at distant locations. In the Real and Hand under conditions attention is focused, since visual and somatosensory stimuli occur at the same location. For body-stimulus related processing, Incongruent and Hand under can be grouped as the visual stimulus does not occur on the subject’s body, while the Real and Two hands conditions can be grouped as the visual stimulus does occur on the participant’s body.

### 2.3 Experimental procedure

In Experiment 1 one block of each condition was administered in pseudorandom order. Each condition included 200 stimulation events. The visuo-tactile stimulus duration was 100 ms and the inter-stimulus interval varied randomly and evenly between 700 ms and 1500 ms. Each block lasted approximately 4 minutes. Experiment 2 contained three blocks of each, the Illusion and Incongruent condition. Each condition included 256 stimulation events. The visuo-tactile stimulus duration was either 100 ms, 125 ms 150 ms or 175 ms. The inter-stimulus interval varied randomly and evenly between 700 ms and 1500 ms. Each block lasted approximately 5 minutes.

In both experiments participants were instructed to use their right hand to press the right arrow key on a computer keyboard when they felt the onset of the illusion and the left arrow key when they lost the feeling of the illusion. Participants sat with their gaze fixed on the LED and wore ear plugs throughout the experiment to reduce the noise caused by the vibration motors.

### 2.4 EEG Recording

Experiments were performed in a darkened and electrically shielded room. EEG signals were continuously recorded using an active 64 channel BioSemi (BioSemi, B.V., The Netherlands) system with Ag-AgCl electrodes mounted on an elastic cap (BioSemi) according to the 10/20 system. Four additional electrodes were placed at the outer canthi and below the eyes to obtain the electro-occulogram (EOG). Electrode offsets were kept below 25 mV. Data were acquired at a sampling rate of 500 Hz using a low pass filter of 208 Hz.

### 2.5 EEG analysis

Data analysis was carried out offline with MATLAB (The MathWorks Inc., Natick, MA) using the FieldTrip toolbox (Oostenveld et al., 2011). Stimulation events and their corresponding triggers were sorted based on condition, presence or absence of the illusion and stimulus length (Experiment 2 only). For the analysis of the Illusion condition only events in which the illusion was present, as indicated by the subjects, were used. For analysis of all other conditions only events in which the illusion was absent were used. EEG data was segmented into epochs of 700 ms (200 ms pre-stimulus to 500 ms post-stimulus) and pre-processed as follows: the data were band-pass filtered between 0.5 Hz and 30 Hz, re-sampled to 200 Hz and subsequently de-noised using independent component analysis (ICA; Debener et al., 2010). In addition, for some subjects highly localized components reflecting muscular artefacts were detected and removed (Hipp and Siegel, 2013; O’Beirne and Patuzzi, 1999). To detect potential artefacts pertaining to remaining blinks or eye movements we computed horizontal, vertical and radial EOG signals following established procedures (Hipp and Siegel, 2013; Keren et al., 2010). We rejected trials on which the peak signal amplitude on any electrode exceeded a level of ±75 μV, or during which potential eye movements were detected based on a threshold of 3 standard deviations above mean of the high-pass filtered EOGs using procedures suggested by Keren et al. (2010). Together these criteria led to the rejection of 34±8 % of trials (mean±SD) in Experiment 1 and of 25±11 % of trials (mean±SD) of trials in Experiment 2. For further analysis the EEG signals were referenced to the common average reference.

Condition averages of the evoked responses (ERPs) and oscillatory power (see below) were computed by randomly sampling the same number of stimulation events from each condition. This was necessary as the number of available trials differed across conditions. Condition averages were obtained by averaging 500 times trial-averages obtained from 80% of the minimally available number of trials.

To analyse oscillatory activity we extracted single trial spectral power for alpha (8-12Hz) and beta (13-25Hz) using a discrete Fourier transformation on sliding Hanning windows with a length of 200 ms. Power values in the range of 100 ms pre-stimulus and 350 ms post-stimulus were averaged across trials. No baseline normalization was performed but within-subject statistical comparisons were used (see below), which make the subtraction of a common baseline unnecessary.

In experiment 1 our primary aims were to determine ERP and oscillatory signatures of the illusion and to compare these to ERP and time-frequency signatures of attentional and body-related processes. While we expected to find significant differences between the Illusion vs. the two control conditions, and in the Attention and Body-stimulus position contrasts, we had no prior expectations about the timing and localisation of significant differences. We hence used spatio-temporal Cluster-based Permutation Analysis to detect significant condition differences (Maris and Oostenveld, 2007). A two-tailed paired t-test was performed for each electrode, and the cluster statistic was defined as the sum of the t-values of all spatially adjacent electrodes exceeding a critical value corresponding to an alpha level of 0.05, and a minimal cluster size of 2. The cluster statistic was compared with the maximum cluster statistic of 2000 random permutations, based on an overall p-value of 0.05. To identify illusion effects we compared Illusion vs Incongruent and Illusion vs Real conditions. For obtaining Body-stimulus position and Attention contrasts we used the four conditions Incongruent, Hand under, Two hands, Real, which differed along the factors of Attention (focussed, divided) and Body-stimulus position (visual stimulus on body, visual stimulus not on body) in a 2x2 design (Fig. 1). To obtain the contrasts for each factor we averaged over the respective conditions belonging to each level and then compared the averages with a cluster permutation test. To calculate the interaction of Attention and Body-stimulus position factors, that is the difference between the differences between the means of one factor, across the levels of the other factor, we subtracted Two hands from Real, and Incongruent from Hand under, and compared these differences with a cluster permutation test.

In experiment 2 our primary aims were to replicate the illusion effect from experiment 1 and to investigate if stimulus duration modulates this effect. For the analysis of evoked responses we selected the time point with the biggest overlap of significant electrodes between Illusion vs Incongruent and Illusion vs Real contrasts as found in experiment 1. We conducted a 2x4 repeated measures ANOVA with the factors illusion presence and stimulus duration on data averaged over the significant electrodes at this time point. For the analysis of oscillatory power we selected the electrodes in the overlap of significant electrodes between Illusion vs Incongruent and Illusion vs Real time-frequency contrasts as found in experiment 1. We conducted a 2x4 repeated measures ANOVA on power in each band. Greenhouse–Geisser correction was applied where sphericity was violated.

## 3 Results

### 3.1 Experiment 1

#### Illusion effect – ERPs

Significant differences (cluster-permutation test, at least p< 0.05) between the Illusion condition and the Incongruent condition emerged around two time points: At 120 ms the Illusion condition showed lower amplitudes in right frontal regions (Tsum = -659.0, p< 0.05) and more positive amplitudes in left parietal areas (Tsum = 490.9, p< 0.05) compared to the Incongruent condition (Fig. 2A). At 330 ms the Illusion condition showed lower amplitudes in frontocentral regions compared to the Incongruent condition and this frontocentral negativity was centred around electrode FCz (Tsum = - 404.4, p< 0.05, Fig. 2A). Significant differences between the Illusion condition and the Real condition emerged around 330ms and were also centred around electrode FCz (Tsum = -823.1, p< 0.05; Fig. 2B). The respective ERPs at electrode FCz suggest that the illusion is characterized by a more pronounced negativity of the evoked activity around 330ms in compared to the two control conditions (Fig. 2C).

**Figure 2.**
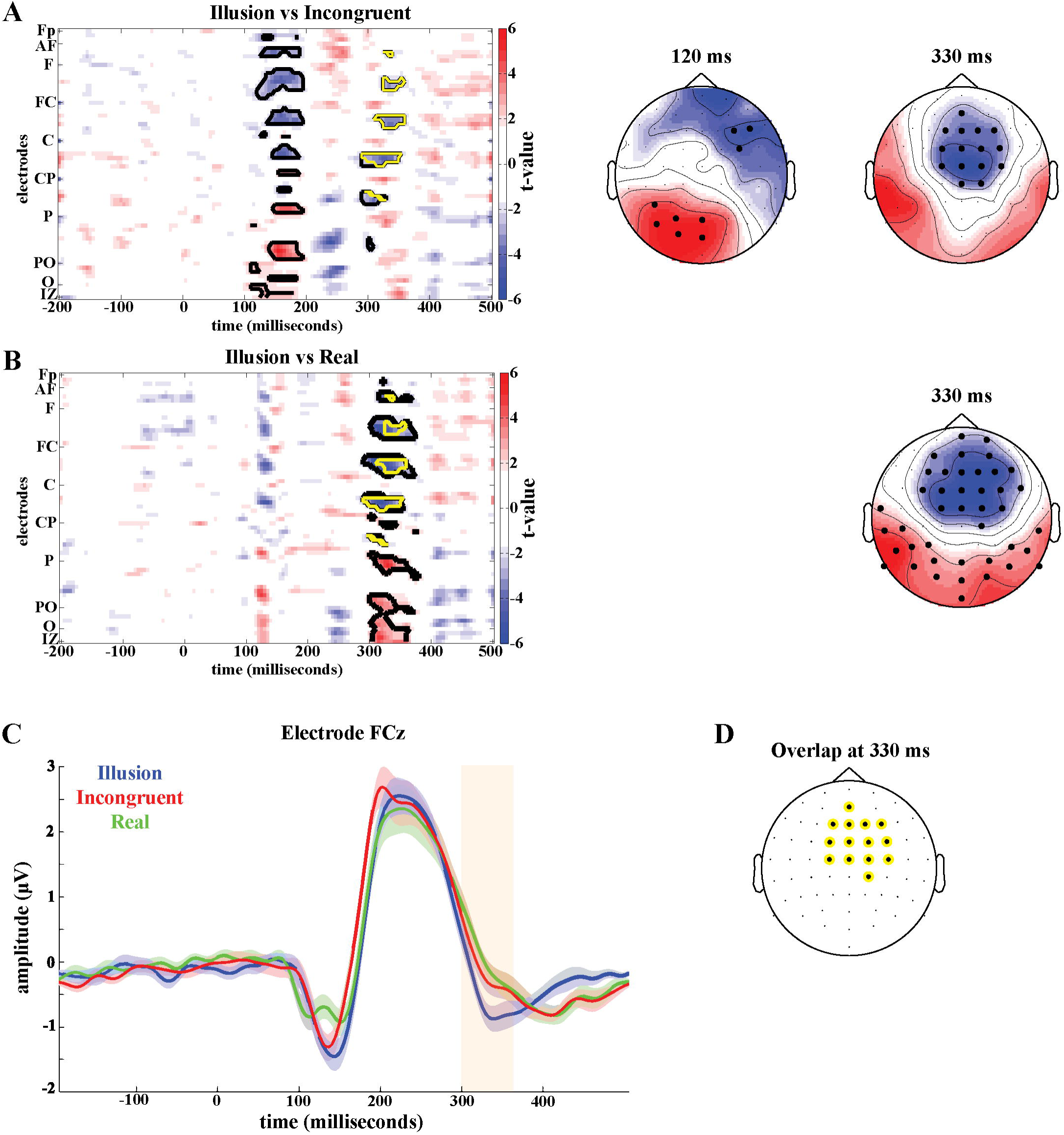
Illusion effect. (A) T-maps of the Illusion vs. Incongruent contrast (top) and the Illusion vs. Real contrast (bottom). Significant clusters (permutation statistics, p < 0.05, n=20) are highlighted in black, significant clusters common to both contrasts are indicated in yellow. (B) Scalp topographies of t-values with significant clusters highlighted. (C) Grand-averaged event-related potentials at FCz of Illusion (blue), Incongruent (red) and Real (green). The shaded areas indicate the standard errors of the mean. (D) Overlap of significant electrodes between Illusion vs. Incongruent contrast and Illusion vs. Real contrast at 330 ms post-stimulus.

To better localize the illusion effect we determined those electrodes that were part of both significant effects around 330 ms, i.e. which were part of the significant time-electrode clusters in the Illusion-Incongruent and Illusion-Real contrasts. The resulting electrodes comprised the medial central and centrofrontal electrodes (Fig. 2D).

#### Illusion effect – Oscillatory activity

The illusion contrasts applied to the power of oscillatory activity revealed significant clusters of 19 electrodes in parietal areas where alpha power (8-12Hz) was lower in the Illusion compared to the Incongruent condition (Tsum = -77.4, p< 0.05; Fig. 3A, top left topography), and lower in the Illusion compared to the Real condition (Tsum = -80.4, p< 0.05, Fig. 3A, bottom left topography). In the beta band (13-25Hz) a cluster of 38 electrodes over frontoparietal regions also showed reduced power during the Illusion condition compared to the Incongruent (Tsum = -109.1, p< 0.05, Fig. 3A, top right topography) and Real conditions (Tsum = -178.2, p< 0.05, Fig. 3A, bottom right topography). The overlap of significant illusion effects for each band is shown in Fig. 3B.

**Figure 3.**
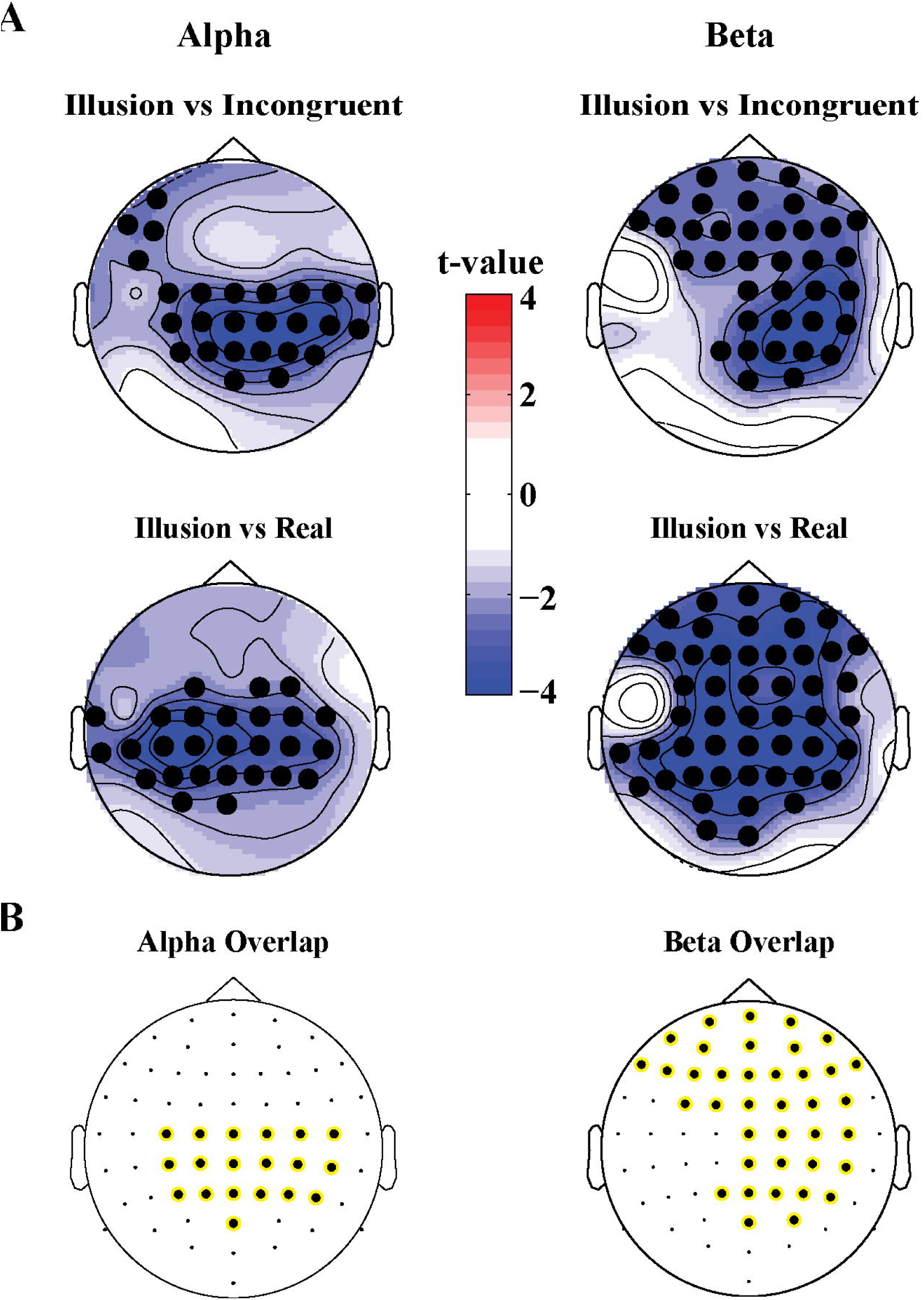
(A) Scalp topographies of t-values for differences in alpha (8-12 Hz, left panel) and beta power (13-25 Hz, right panel) for the Illusion vs Incongruent (top panel) and Illusion vs Real (bottom panel) contrast. Significant clusters (permutation statistics; p < .05, n=20) are highlighted in black. (B) Overlap of significant clusters between the Illusion vs Incongruent and Illusion vs Real contrasts.

#### Attention and Body-stimulus position contrasts

Potential confounding effects of changes in spatial attention and body-stimulus position were quantified using four additional experimental conditions analysed in a 2x2 design (Fig. 1). In the time domain no significant effects were found when analysing the interaction between the factors Attention and Body-stimulus position. However, significant effects emerged in the attention contrast around 100 ms (Positive cluster: Tsum = 701.0, p< 0.05; Negative cluster: Tsum = 728.0, p< 0.05) and 250 ms (Positive cluster: Tsum = 687.7, p< 0.05; Negative cluster: Tsum = -470.4, p< 0.05; Fig. 4A) in frontal and parietal regions. Significant effects in the body-stimulus position contrast emerged around 180 ms centred around electrode FCz (Tsum = -474.6, p< 0.05; Fig. 4B).

While the timing and location of the attention effects do not resemble the illusion effect, the topography of significant effects in the body-stimulus position contrast closely resembles the topography of the illusion effect (c.f. Fig. 2D). The electrodes consistently involved in both effects comprised medial central and centrofrontal electrodes (Fig. 4C), making it possible that potentially similar regions are involved in mediating the illusion and body-stimulus effects, but reflect these at distinct latencies relative to the stimulus.

**Figure 4.**
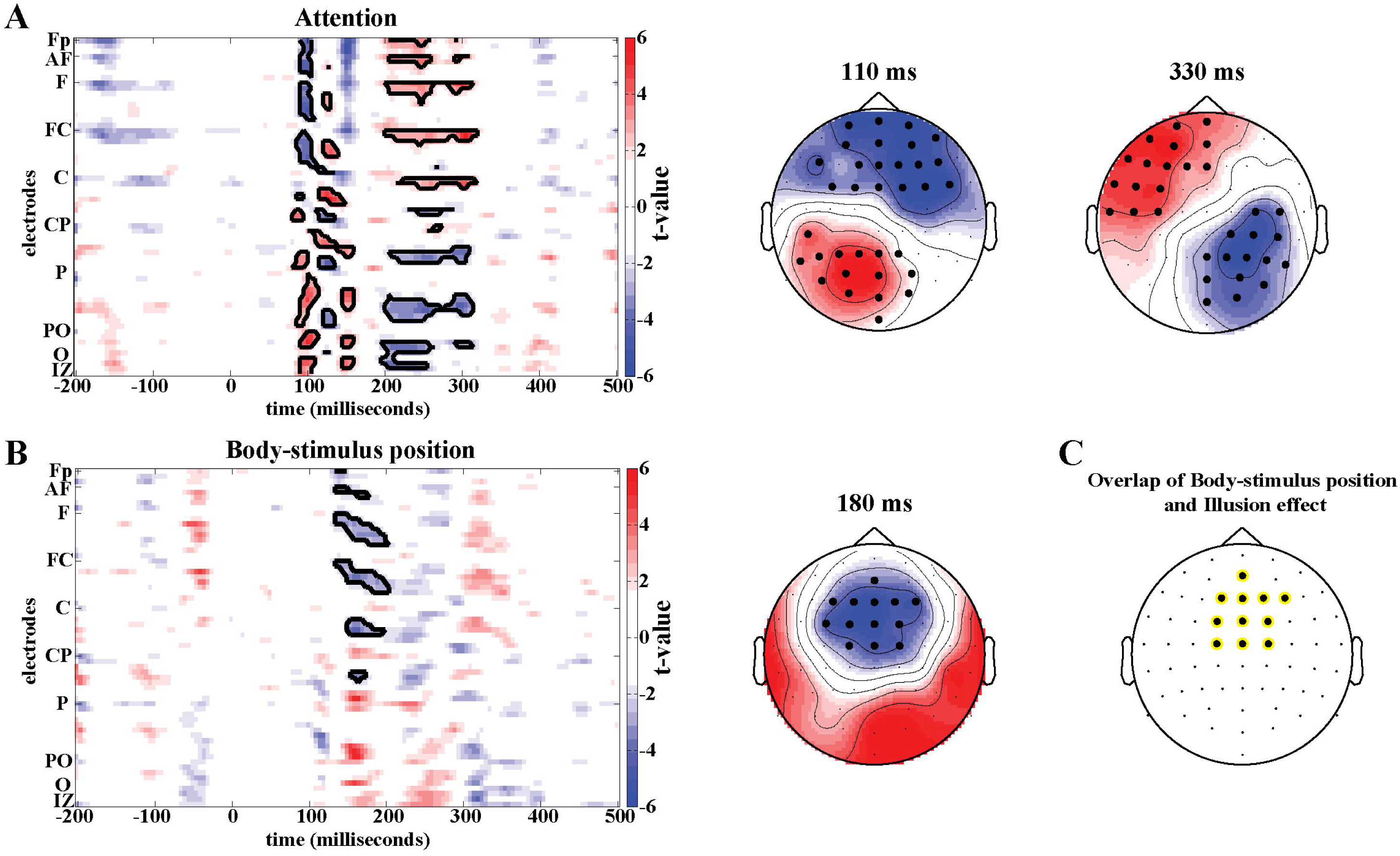
Contrasts for the effects of Attention and Body-stimulus position. (A) T-maps for Attention (top) and Body-stimulus position contrasts (bottom). Significant clusters (permutation statistics; p < 0.05, n=20) are highlighted in black. (B) Scalp topographies of t-values with significant clusters highlighted. (C) Overlap of significant clusters between the illusion effect (from Fig. 2D) and the body-stimulus position effect.

We found no significant differences in oscillatory responses in the attention and body-stimulus position contrasts in either the alpha (8-12Hz) or beta band (13-25 Hz).

### 3.2 Experiment 2

#### Illusion effect – ERPs

In the second experiment we compared the Illusion to the Incongruent condition while manipulating the duration of the visuo-tactile stimulation. We then performed a repeated-measures ANOVA on the ERP amplitudes at the time-electrode cluster identified by the illusion effect in experiment 1 (c.f. Fig. 2D) to test the effects of illusion and stimulus duration (Table 1). This confirmed a main effect of illusion at 330ms in this second dataset (F_(1,19)_=16.08, p< 0.05, η^2p^ =0.46), and revealed an effect of stimulus duration (F_(1.63,31.02_)=21.318, p< 0.05, η^2p^ =0.53) but no significant interaction (F_(2.81,53.40_)=0.235, p=0.860, η^2p^ =0.012).

**Table 1.** Group means and standard deviations of amplitudes (μV) at 330ms post-stimulus in experiment 2.

| Condition | Stimulus duration |  |  |  |
| --- | --- | --- | --- | --- |
|  | 100ms | 125ms | 150ms | 175ms |
| **Illusion** | -0.6652 (0.8934) | -0.3829 (0.6558) | -0.0917 (0.5668) | 0.0138 (0.4904) |
| **Incongruent** | -0.4447 (0.7040) | -0.1790 (0.7211) | 0.2105 (0.6217) | 0.2503 (0.6161) |

#### Illusion effect – Oscillatory activity

For alpha power we found a main effect of illusion (F_(1.00,19.00)_=16.407, p< 0.05, η^2p^ =0.46) but no effect of stimulus duration (F_(2.69,51.08)_=2.822, p=0.053, η^2p^ =0.13) and no significant interaction (F_(2.36,44.85)_=2.860, p=0.059, η^2p^ =0.13). For beta power we found a main effect of illusion (F_(1.00,19.00)_=15.337, p< 0.05, η^2p^ =0.45) but no main effect of stimulus duration (F_(2.36,44.84)_=2.917, p=0.056, η^2p^ =0.13). However, a significant interaction between illusion presence and stimulus duration was present (F_(2.28,43.33)_= 7.533, p< 0.05, η^2p^ =0.28). This interaction appeared to be driven by higher beta power for the stimulus duration of 100ms compared to the other durations in the illusion condition (Table 2).

**Table 2.** Mean values and standard deviations of oscillatory power in alpha (8-12 Hz) and beta band (13-25 Hz) in experiment 2.

| Frequency | Condition | Stimulus duration |  |  |  |
| --- | --- | --- | --- | --- | --- |
|  |  | 100ms | 125ms | 150ms | 175ms |
| **Alpha**<br>**(8-12Hz)** | **Illusion** | 3.9137 (1.9996) | 3.5172 (1.7485) | 3.6505 (1.7688) | 3.6360 (2.0241) |
|  | **Incongruent** | 4.3655 (2.1172) | 4.3723 ( 2.0797) | 4.4066 (2.1775) | 4.3273 (2.0935) |
| **Beta**<br>**(13-25Hz)** | **Illusion** | 1.0390 (0.4441) | 0.9808 (0.4092) | 0.9843 (0.4028) | 0.9748 (0.4180) |
|  | **Incongruent** | 1.0778 (0.4461) | 1.1071 (0.4672) | 1.0895 (0.4380) | 1.0897 (0.4432) |

## 4 Discussion

We studied the neurophysiological correlates of the rubber hand illusion using a fully automated and precisely-timed visuo-tactile setup and a combination of experimental conditions. Across two studies and two control conditions we reliably found an illusion-related attenuation of ERPs around 330ms over frontocentral electrodes. This effect was not related to attention or body-stimulus position confounds and was robust against changes in stimulus duration. We furthermore found that oscillatory activity in the alpha and beta bands was reliably reduced during the illusion. We thereby provide multiple neural markers of the RHI.

### Illusion effects in evoked responses

Several previous EEG studies have aimed to understand the neural correlates and mechanisms underlying the illusory percept of body ownership in the RHI. These studies compared the evoked responses associated with the tactile stimulus on the participant’s hand between conditions inducing the illusion and control conditions. The rationale behind this approach is to see whether and how the cortical representation of the tactile stimulus changes when its subjective location changes from the actual hand to the rubber hand. Previous studies differed regarding the latency of such an illusion-correlate in ERPs, reporting either early effects around 55 ms (Zeller et al., 2015) or much later effects around 460 ms (Peled et al., 2003). However, both studies relied on the manual stimulation by a brush handled by an experimenter, whereby each individual brush stroke can differ in timing and intensity. This variability in the sensory stimulus can be detrimental for measuring the timing and shape of the respective sensory evoked responses. To overcome this problem we here designed an automated setup that allows visuo-tactile stimulation with great temporal fidelity and consistency across trials. Furthermore, we asked subjects to indicate the onset of the rubber hand illusion during each trial and hence were able to include only those stimulation events in the analysis during which subjects actually reported the presence of the RHI. To facilitate this we only considered participants that had previously and reliably experienced ownership over a rubber hand and were familiar with the sensations associated with onset and presence of the RHI as determined by a pilot session.

To establish neural correlates of the RHI a comparison of the illusion condition with a control condition is required. Most previous ERP studies relied on the Incongruent condition in which the rubber hand is placed at an anatomically incongruent angle, or relied on the Real condition in which the rubber hand is absent and stimulation occurs on the real hand in view (Peled et al., 2003; Zeller et al., 2015, 2016). Using only one control condition makes the implicit and critical assumption that the illusion and control conditions differ only in a single factor, the presence of the subjective illusion. Yet, closer inspection of these conditions suggests that these may differ by other factors as well, such as focus of attention and body-stimulus position in the Incongruent condition, or the absence of a rubber hand in the Real condition. We therefore relied on the combination of control conditions to identify potential changes in evoked activity that are reliably associated with the illusion. The need to consider multiple control conditions is further demonstrated by the observation that some significant ERP effects were observed only in one of the two contrasts (c.f. Fig. 2). For example, the Illusion-Incongruent difference revealed a significant effect around 150 ms, which was absent in the Illusion-Real difference, and hence unlikely is a correlate of the subjective illusion. This suggests that results on the neural correlates of illusory body ownership that were obtained using a single control condition have to be considered with care.

We found neural activations that were reliably associated with the illusion only at longer latencies (here 330ms) over frontocentral regions. Furthermore, this illusion effect did not interact with changes in stimulus duration. Together this suggests that these activations do not reflect processes related to early sensory encoding but rather reflect late and higher-order processes. Thereby our results differ from Zeller et al. (2015) who reported illusion related activity as early as 55 ms, but also differ from those of Peled et al. (2003), who found illusion related activity around 460ms. The discrepancies are possibly due to several factors: First, these previous studies relied on the manual stimulation by a brush, as opposed to the automated visual-tactile stimuli in the current study. Second, Zeller et al. restricted their analysis to activity before 300 ms post-stimulus, while Peled et al. only tested at specific time points not including 330 ms. This makes it difficult to compare significant effects across studies, as each relied on distinct time windows where potential effects were expected and contrasted using methods for multiple comparison. Third, the study by Zeller et al. relied on a rather small sample size (n=13), while we here relied on a sample size of n=20 participants in each experiment, which is considered to be the minimal sample size for neuroimaging studies based on concerns of reporting false positive results (Poldrack et al., 2017; Simmons et al., 2011). Fourth, the study of Zeller et al. reported significant illusion effects only for stimulation on the right hand, while we here focused on the subject’s left hand, as previous studies have suggested a strong link between the right hemisphere and awareness of the subjective experience of body ownership (Frassinetti et al., 2008; Karnath and Baier, 2010; Tsakiris et al., 2007). Last but not least, we replicated the illusion effect around 330 ms in two independent studies, providing further evidence for the robustness of our result.

### Neural origins of illusion-related activations

The frontocentral location of the illusion effect in the current study provides support for a pivotal role of premotor and possibly intraparietal areas in illusory hand ownership. Several fMRI studies have consistently associated activity in the ventral premotor cortex and posterior parietal with the illusory percept of ownership and hand position in the RHI (Brozzoli et al., 2012; Guterstam et al., 2015; Limanowski and Blankenburg, 2015; Petkova et al., 2011). Furthermore, Limanowski et al. (2015) and Guterstam et al. (2015) reported increased functional coupling between intra-parietal regions and premotor cortices during the illusion compared to control conditions. Both regions are ideal candidates for mediating the multisensory integrative processes that underlie the RHI. They process signals involved in self-attribution of the hand (Ehrsson, 2004; Evans and Blanke, 2013; Tsakiris et al., 2007) and analogous regions in the monkey brain have been found to contain trimodal neurons that integrate tactile, visual and proprioceptive signals (Fogassi et al., 1996; Graziano et al., 1997; Graziano and Gandhi, 2000; Iriki et al., 1996). Based on the topography of illusion-related ERP effects our data further corroborate a central role of motor-related regions in the body illusion.

This interpretation is further supported by the timing of the illusion effect, which matches results from intracranial recording studies, which have reported correlates of multisensory integration between 280 and 330 ms over precentral and postcentral regions adjacent to premotor cortex and IPS (Quinn et al., 2014). Similar late latencies were also reported for the integration of visual and somatosensory in parietal association cortex (Lippert et al., 2013). The attenuation of the evoked potential at 330ms during the illusion condition observed here could thus be indicative of the integration of visual, tactile, and proprioceptive information within the parietal-premotor network, which then results in the illusory percept of ownership and hand position in the RHI.

We did not administer any behavioural or physiological measures to measure the RHI, such as proprioceptive drift measurements or changes in body temperature. The reason for this was twofold. Firstly, we relied on a subjective measure of the illusion, as it allowed for uninterrupted recording of EEG data across all conditions. Secondly, our study aimed to identify the correlates of the ownership aspect of the RHI. As shown recently, proprioceptive drift does not provide a reliable assessment this ownership aspect (Rohde et al., 2011). Rather, subjective ownership and the proprioceptive drift can be dissociated, with the latter measuring the spatial updating of the body in space rather than the strength of ownership over the rubber hand itself.

### Illusion, attention and body-stimulus position

We used additional control conditions to reliably dissociate the neural correlates of the RHI from attention and body-stimulus position related activity. Specifically, we identified the timing and location of attention / body-stimulus position related effects and compared these to the activations revealed by the two statistical contrasts obtained from the Illusion. By comparing conditions where the visual stimulus was near the body with conditions where the visual stimulus was far from the body, we found body-stimulus position related processing to be associated with activity in frontocentral areas around 180 ms. This is in line with previous studies investigating the influence of proximity of hands and visual stimuli. For example, Reed et al. (2013) recorded ERPs during a visual detection task in which the hand was placed near or kept far from the stimuli. Similar to the results of the current study, they found increased negativity in the Nd1 component around 180 ms in the near hand condition (see also Sambo and Forster, 2009). The timing of the body-stimulus position related activity (∼180 ms) was notably different from that of the illusion effect (∼330 ms). This differentiates the illusion effect from body-stimulus position related activity. However, the topography of the body-stimulus position related activity at 180 ms was highly similar to that of our illusion effect at 330 ms. Thus, it is possible that both effects may emanate from the same cortical networks related to body processing. Support for this comes from a study by Brozzoli et al. (2012) who measured BOLD response while presenting participants with visual stimuli occurring next or distant from their hands. Their results indicated increased activity in premotor and intraparietal cortices in the condition where the stimulus was close to the hand compared to the condition where the stimulus was distant form the hand. Similar results were obtained when the participant’s hand was replaced by a rubber hand on which the RHI was induced (Brozzoli et al., 2012). This suggests that both, the effects of body-stimulus position and the illusion may originate from processing in the intraparietal-premotor network but do so at different latencies relative to stimulus onset, further corroborating that the ERP correlates of the illusory percept reflect sensory integration processes in the parietal-premotor network.

We found attention related activity in frontal and parietal regions around 100 ms and around 250 ms. This timing is in agreement with previous ERP studies on visual-tactile attention which presented simultaneous stimuli in close proximity or at distant locations (Eimer and Driver, 2001; Sambo and Forster, 2009) and reported modulations of amplitudes between 80-125 ms and 200-280 ms associated with the induced changes in spatial attention. Interestingly the timing and location of activity related to attentional processing in our study is similar to the timing and location of early differences between Illusion and Incongruent. This could mean that these early differences in evoked potentials between Illusion and Incongruent condition are not directly related to the illusion but rather reflect the difference in attention focus between the two conditions. This underlines that the Incongruent condition, one of the most commonly used control condition in EEG experiments on the RHI, should be used with caution when trying to determine the neurophysiological correlates of the RHI.

### Illusion effects in oscillatory activity

The analysis of oscillatory activity revealed that illusory hand ownership resulted in a relative decrease of oscillatory power in the alpha and beta bands. Modulations of alpha power have previously been implicated in the rubber hand illusion (Evans and Blanke, 2013) as well as the full body illusion (Lenggenhager et al., 2011). Our results are also in good agreement with those from a recent study on the somatic RHI (Faivre et al., 2017), a variant of the conventional RHI. Very similar to our results this study found a relative decrease in alpha power over frontocentral regions contralateral to stimulation site and a relative decrease in beta power bilaterally over frontoparietal regions during the illusion. Combined with the consistency of these power decreases across two control conditions and two experiments as shown here, this implicates that the decrease in alpha and beta power during the illusion is not associated with visual information or a specific control condition. Instead, it is likely to be directly tied to the feeling of ownership during the illusion itself, and hence constitutes a robust physiological marker of body ownership.

### Conclusion

We identified neurophysiological correlates of the rubber hand illusion in a reduction of alpha and beta power as well as in an attenuation of evoked responses around 330 ms over central electrodes. The attenuation of evoked responses is likely to reflect the integration of visual, somatosensory and proprioceptive information during the illusion, which then leads to the experience of ownership over the rubber hand. Our results furthermore emphasize the need to consider multiple control conditions in studies on body illusions, to avoid misidentifying effects related to changes in body-stimulus position or attention for correlates of illusory body ownership.

## 5 Acknowledgements

This work was supported by the European Research Council (to C.K. ERC-2014-CoG; grant No 646657) and a studentship from the UK Economic and Social Research Council ESRC (to I.R., award No 1612065).

### 6 Author Contributions

IR and CK designed the study, IR ran the experiments, IR and CK analysed the data and wrote the manuscript.

